# Dopamine is a double-edged sword: Enhancing memory retrieval performance at the expense of metacognition

**DOI:** 10.1101/274159

**Authors:** Mareike Clos, Nico Bunzeck, Tobias Sommer

## Abstract

While memory encoding and consolidation processes have been linked with dopaminergic signaling for a long time, the role of dopamine in episodic memory retrieval remained mostly unexplored. Based on previous observations of striatal activity during memory retrieval, we used pharmacological fMRI to investigate the effects of dopamine on retrieval performance and metacognitive memory confidence in healthy humans. Dopaminergic modulation by the D2 antagonist haloperidol administered acutely during the retrieval phase improved recognition accuracy of previously learned pictures significantly and was associated with increased activity in the SN/VTA, locus coeruleus, hippocampus and amygdala during retrieval. In contrast, confidence for new-decisions was impaired by unsystematically increased activity of the striatum across confidence levels and restricted range of responsiveness in frontostriatal networks under haloperidol. These findings offer new insights into the mechanisms underlying memory retrieval and metacognition and provide a broader perspective on the presence of memory problems in dopamine-related diseases and the treatment of memory disorders.

## Introduction

Numerous studies have specified the contribution of the medial temporal lobe as well as prefrontal and parietal areas to the retrieval of episodic information. However, besides this established retrieval network also striatal activity is consistently observed during memory retrieval (Kim, 2013), which might point to a hitherto uncharted role of dopaminergic neurotransmission in retrieval processes. In contrast to retrieval, encoding and consolidation of memories are well-known to require the stimulation of dopamine receptors in the hippocampus, striatum, amygdala and prefrontal cortex (PFC) (Mele *et al*, 2004; Rocchetti *et al*, 2015; Rossato *et al*, 2013) as part of a functional loop that orchestrates the formation of new memories (Axmacher *et al*, 2010; Lisman and Grace, 2005). Striatal dopaminergic influence during retrieval, on the other hand, has been suggested to reflect motivational aspects such as higher subjective value of successfully retrieving old than rejecting new items in a memory test (Han *et al*, 2010) or higher memory confidence during retrieval (Schwarze *et al*, 2013). Intriguingly, however, dopamine might also affect the actual retrieval of episodic information, for example it could potentially support cognitive control processes during retrieval (Scimeca and Badre, 2012) or improve the signal-to-noise ratio of memory representations (Frank, 2005; Warren *et al*, 2016; Yousif *et al*, 2016). In a previous fMRI study involving a picture recognition task, we provided evidence for independent retrieval-related and memory confidence signals in overlapping striatal regions (Clos *et al*, 2015). We here investigated directly whether these striatally-mediated memory processes are indeed dopaminergic. In order to disentangle the potential dopaminergic effects on memory confidence and retrieval performance, we designed a randomized, placebo-controlled between-group pharmacological fMRI study. We used the same recognition task as in our previous study (Clos *et al*, 2015), in which the participants had to rate the old/new status of pictures and their memory confidence on a combined 6-point confidence scale. Dopamine level was manipulated during retrieval by 2 mg of the D2-antagonist haloperidol administered only after encoding.

## Materials and Methods

### Participants and group allocation

Fifty-four volunteers (14 males, mean age 24±2.9 years) participated in the study. We used a double-blind, placebo-controlled between-group design, assigning 27 participants randomly to either placebo or haloperidol group. The sample size was based on our experience in earlier fMRI studies using the same picture recognition paradigm (Clos *et al*, 2015; Schwarze *et al*, 2013) and previous pharmacological fMRI studies comparing haloperidol and placebo using between-group designs (Cole *et al*, 2013; Menon *et al*, 2007; Oei *et al*, 2012; Pessiglione *et al*, 2006; Pleger *et al*, 2009; Wrobel *et al*, 2014). We chose for a between-group design rather than for the more powerful within-group crossover design to avoid interference between the highly similar picture stimuli used in the recognition paradigm and because of additional tasks measured in this sample (Clos *et al*, in revision), which were incompatible with a within-group design. The random group assignment was conducted by TS, who did not interact with the participants at any time. All other experimenters were blind with regard to group assignment and group allocation was only revealed after the measurement of the last participant was completed. The study was approved by the local ethics committee of the Hamburg medical association and written consent was given by each participant prior to the start of the study. Additionally, all participants were screened by a physician for previous or current physical or mental diseases, medication or drug use, ensuring that only healthy participants were included in the study. The participants were informed about the purpose and the course of the study and about the potential risks and side effects of haloperidol. Moreover, the participants were instructed to restrain from caffeine, nicotine, and alcohol on the day of the fMRI testing. Repeated blood pressure and pulse measurements together with questionnaires on adverse medication effects and current mood ensured the well-being of the participants after haloperidol/placebo administration. All participants were asked to indicate the substance (haloperidol or placebo) they thought they had received and how certain they were of this guess at the end of the study.

### Experimental design

#### Behavioral working-memory baseline and encoding session (T0)

Encoding for the fMRI experiment took place outside the scanner on the first day within a one-hour baseline testing session (no medication given). The participants first completed working memory (WM) baseline tests on a computer. We measured their performance on a self-paced complex span task (Unsworth *et al*, 2009), where the location of 2-5 squares within a 16-square pattern had to be remembered while simultaneously judging the symmetry of abstract pictures (symmetric/asymmetric). Subsequently, the participants performed a self-paced digit and block span task (Kessels *et al*, 2008). Digit sequences presented auditorily via headphones had to be entered via the keyboard after presentation of the last digit in either forward or backward order. For the forward digit span, the sequence increased from 3 digits to maximally 8 digits (depending on performance). For the backward digit span, the sequence increased from 2 digit to maximally 7 digits. Similarly, during trials of the block span task, the participants saw a pattern of objects, whose location and (forward/backward) appearance order they had subsequently to indicate via mouse clicks on these objects. For the forward block span, the sequence increased from 2 objects to maximally 8 objects. For the backward block span, the sequence increased from 2 to maximally 7 objects.

For the recognition memory task, we used the same encoding paradigm as previously (Clos *et al*, 2015). In total 80 unfamiliar photos of outdoor scenes (Peelen *et al*, 2009) were presented for 800 ms. The participants had to indicate the category for each picture (i.e. whether it contained cars or people) by pressing one of two buttons on the keyboard (Fig. 1 A). Each picture was followed by an active baseline condition (ISI 8-12 s, arrow pointing task) in which the direction of arrows presented for 800 ms had to be indicated by button press.

**Figure 1:**
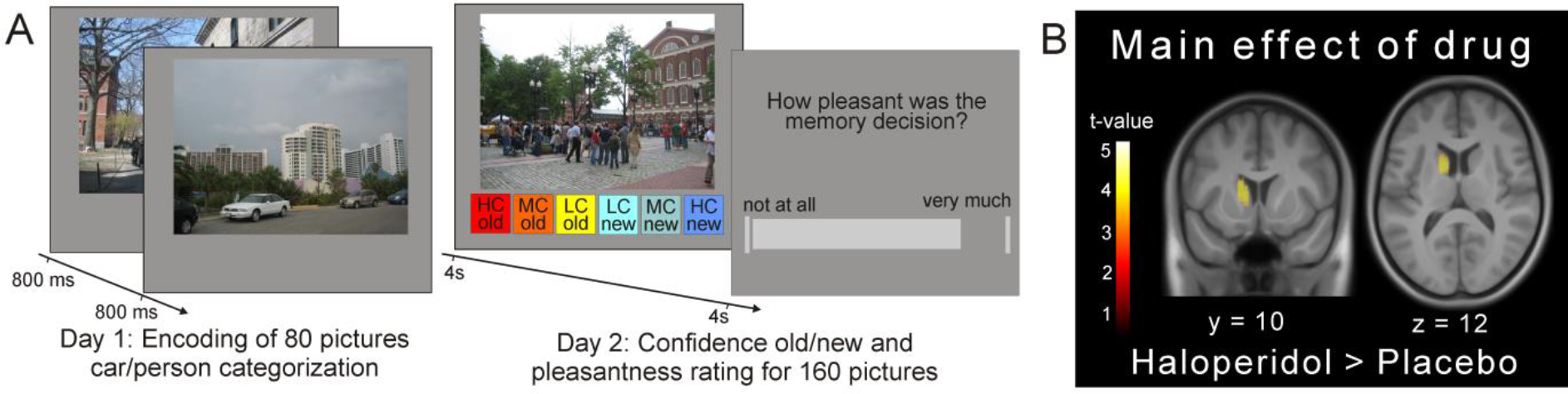
Task design and general drug effects. A) Participants encoded pictures of outdoor scenes outside the scanner. Picture recognition took place under haloperidol/placebo in the fMRI scanner on the next day. B) Haloperidol effect across all trials on brain activity compared to placebo. Note that increased activity in the right dorsal striatum was also present but its extent (k = 76 voxels) and height (z = 3.52) did not pass the FWE-corrected threshold of p <. 05. The inverse contrast (placebo > haloperidol) revealed no significant activity. Activation maps are thresholded at p <. 05 (FWE-corrected at cluster-level using a cluster forming threshold at voxel level of p <. 001). HC = high confidence, MC = medium confidence, LC = low confidence.

#### Recognition session under haloperidol/placebo in the fMRI scanner (T1)

On the next day, the participants filled out questionnaires assessing their current mood and potential adverse effects of the medication. Blood pressure and pulse were controlled by a physician prior to drug administration. Next, the participants received a tablet containing either 2 mg haloperidol or placebo (based on prior random assignment). In the following waiting period of 2.5 hours, blood pressure and pulse were checked again 30 minutes and 2 hours after drug administration. Also the questionnaires on current mood and adverse medication effects were filled out twice again by the participants. After a brief practice round and familiarization with the recognition task, the participants started the picture recognition inside the scanner approximately 2.5 hours after tablet ingestion (Fig. 1A). During the recognition task, all 80 previously encoded pictures were presented randomly intermixed with 80 new pictures (lures). For each picture (presented for 4 s, ISI 2-4 s), participants indicated the old/new status and their subjective memory confidence on a combined 6-point confidence scale (1 - “high confidence old”, 6 - “high confidence new”) by pressing one of six buttons. The boxes corresponding to the various levels of confidence old/new were horizontally arranged in each trial in random order to minimize transfer-effects to the next rating. Feedback about the correctness of the response was not given. 2-4 seconds after each confidence old/new rating, participants indicated how pleasant the preceding memory retrieval was on a visual analogue scale (VAS) ranging from 1 (“not at all”) to 100 (“very much”). The pleasantness rating was measured on the continuous VAS (rather than on a categorical scale as the confidence rating) to minimize an implicit transfer from the confidence to the pleasantness ratings. The completion of this task took approximately 45 minutes. After the scanning session, the participants completed another round of the mood and adverse effect questionnaires and their blood pressure and pulse was checked a last time. Another behavioral testing session on the computer followed, including again the working memory tasks (complex span, block span and digit span).

#### Image acquisition and pre-processing

Functional MR images were obtained during recognition on a 3T system Siemens Trio using single-shot echo-planar imaging with parallel imaging (GRAPPA (Griswold *et al*, 2002), in-plane acceleration factor 2) and simultaneous multi-slice acquisitions (Feinberg *et al*, 2010; Moeller *et al*, 2010; Setsompop *et al*, 2012; Xu *et al*, 2013) (“multiband”, slice acceleration factor 2; TR = 1.98s, TE = 26ms, flip angle = 70°, 64 axial slices, voxel size 2×2×2 mm^3^). The corresponding image reconstruction algorithm was provided by the University of Minnesota Center for Magnetic Resonance Research. In addition, an anatomical high-resolution T1-weighted image (TR = 2.3s, TE = 2.98ms, flip angle = 9°, 192 sagittal slices, voxel size 1×1×1 mm^3^) and an anatomical magnetization transfer (MT) image (TR = 14ms, TE = 3.2ms, flip angle = 6°, 240 coronal slices, voxel size 1×1×1 mm^3^) was acquired for each participant. The data were pre-processed using SPM12 (http://www.fil.ion.ucl.ac.uk/spm/). The first five EPI images were discarded to allow for magnetic-field saturation. EPI images were corrected for motion and for the interaction between motion and distortion (using the unwarping procedure). Anatomical T1-weighted images were normalized to standard MNI space using DARTEL normalization. Subsequently, the EPI images and the MT image were co-registered with the normalized T1 image and the DARTEL normalization parameters were applied to the EPI images and the MT image. Finally, these normalized EPI images were spatially smoothed with a Gaussian kernel of 8 mm full width at half- maximum.

### Data analysis

#### Behavioral data

All behavioral data were analyzed using SPSS 21.0.0 (SPSS; Chicago, IL) and Matlab R2013a (The MathWorks; Natick, MA). Post-hoc tests were Bonferroni-corrected where appropriate and corrected p-values are reported.

#### Side effects and mood

The scores on the adverse medication effects questionnaire were summed together per measurement time point and analyzed relative to baseline for group differences using a repeated measures ANOVA with the factors group and time. Similarly, pulse and blood pressure measurements were analyzed relative to baseline for group differences using a repeated measures ANOVA with the factors group and time. The 16-item mood questionnaire was analyzed by reversing inverted items and log-transforming all scales before grouping the items into the three dimensions “alertness”, “calmness” and “contentment” (Bond and Lader, 1974). The resulting three dimensions were compared between the groups relative to baseline using 2×2 repeated measures ANOVAs with the factors group and time.

#### Recognition memory task

For the recognition performance at T1, the responses on the 6-point confidence scale were used to compute the accuracy of each trial based on the factual old- or new-status of the item. Trials were sorted post-hoc into the response categories hits (correct old-responses), correct rejections (CR, correct new-responses), false alarms (FA, incorrect old-responses) and misses (incorrect new- responses). Discrimination sensitivity (d’) and response bias (c) were computed according to the signal detection approach (Macmillan and Creelman, 2004) both across all confidence levels and separately for each confidence level. Discrimination sensitivity was calculated as *d’* = *z(H)* – *z(FA)*, where z represents the inverse of the cumulative normal distribution and *H* = *p*(response = old |stimulus = old) and *FA* = *p*(response = old | stimulus = new). Response bias was calculated as *c* = -0.5[*z(H)* + *z(F)*]. The resulting d’ and c parameters were compared between groups using independent samples t-tests and d’ and c parameters per confidence level were analyzed with 2×3 repeated-measures analyses of variance (ANOVAs) with the factors group and confidence level. We moreover analyzed the effects of haloperidol on recollection vs. familiarity processes, which are thought to differentially contribute to memory retrieval (Yonelinas, 2002). According to the dual-process model of recognition memory (Yonelinas *et al*, 2010), recollection is a threshold process which constitutes the recall of detailed and specific qualitative information. In contrast, familiarity is a quantitative signal detection process, which reflects more global aspects of memory strength. We used the dual process signal detection (DPSD) model as implemented the ROC Toolbox (Koen *et al*, 2016) to estimate the parameters of recollection and familiarity as well as the receiver operating characteristic (ROC) curves for each participant based on the individual confidence ratings. The resulting recollection and familiarity parameters were averaged and compared between groups using independent-samples t-tests.

Metamemory for old and new picture trials was quantified using the meta-d’ framework (Maniscalco and Lau, 2012). Meta-d’ aims to quantify metacognitive sensitivity, that is, how well the observer’s confidence ratings discriminate between correct and incorrect responses. Meta-d’ can be evaluated with respect to d’ in order to take into account the observer’s discrimination sensitivity by calculating meta-d’ – d’ (meta-d’ difference) as meta-d’ is expressed in the same units as d’. Suboptimal metacognition is then reflected by meta-d’ values below zero (meta-d’ < d’) and enhanced metacognition is reflected by meta-d’ values above zero (meta-d’ > d’) (Maniscalco and Lau, 2012). Response-specific meta-d’ values per trial type (old/new) (Maniscalco and Lau, 2014) were calculated using the MATLAB code available at http://www.columbia.edu/∼bsm2105/type2sdt, averaged and compared between groups using a 2×2 repeated-measures ANOVA with the factors group and old/new. For one placebo participant, the modeled meta-d’ for old picture trials was more than 4 standard deviations above the mean (z-score > 4.4) and thus this participant was excluded from the group analysis.

Moreover, mean confidence levels (coded from 1 to 3) were computed per response category (hits, CR, FA, misses) and compared between groups using a 2×2×2 repeated-measures ANOVA with the factors group, old/new rating and accuracy. Differences in the frequency of low, medium and high confidence responses were analyzed using a 2×2×2×3 repeated-measures ANOVA with the factors group, old/new rating, accuracy and confidence level to test whether the confidence responses were selected equally often in both groups. Reaction times (RT) differences as well as pleasantness ratings could not be analyzed by the full 2×2×2×3 model due to the scarceness of trials with high confidence incorrect responses. Therefore, we analyzed RT (based on the median RT per participant) and pleasantness data using an independent t-test to test for group differences and multiple repeated-measures ANOVAs to test for all possible group×condition interactions (2×2×3 model collapsed across accuracy with factors group, old/new rating and confidence; 2×2×3 model collapsed across old/new rating with factors group, accuracy and confidence; 2×2×2 model collapsed across confidence with factors group, accuracy and old/new rating).

#### WM

Due to fatigue two haloperidol participants did not complete the post-fMRI T1 WM span tasks. Additionally, partial data loss due to technical problems affected three haloperidol and one placebo participant in the T1 WM span tasks. Working memory scores were computed for each of the WM span tests (complex span performance, block span performance forward/backward, digit span performance forward/backward) as well as for the accuracy and the RT of the symmetry rating of the complex span test. The individual T0 baseline WM measures acquired prior to drug administration were compared between groups using independent samples t-tests. We summarized the five WM span scores into a single T0 and T1 WM span summary score using two different approaches. Firstly, z-scores of each WM span test were computed and averaged into a T0 baseline score and T1 score, respectively. Secondly, for T0 and T1 data separately, a principal components analysis (PCA) was conducted on the individual WM span scores and the resulting first component of T0 and of T1 data was used as a WM span summary score. Group differences were again examined by means of an independent t-test (T0 baseline WM span summary score) and a repeated measures ANOVA with the factors group and time to test for changes in WM from pre- to post-drug administration. The T0 baseline WM span summary score was moreover used as a covariate in the behavioral and fMRI analysis of memory and confidence effects to control for the potential influence of the individual dopamine baseline on medication response (Cools *et al*, 2008).

#### Imaging data

One placebo dataset was acquired only behaviorally due to scanner problems on that day. For the remaining 53 datasets, univariate single subject (first-level) and group (second-level) statistics were conducted using the general linear model as implemented in SPM12. On the first level, delta functions marking trial-onsets of all corresponding events were convolved with the canonical hemodynamic response function to create an event-related regressor for each condition. Low-frequency signal drifts were removed by employing a highpass filter with a cut-off period of 128 s.

First, in order to test for general group differences throughout the recognition memory task, we set up a first-level model containing a regressor representing all picture onset trials. The continuous variable pleasantness was modeled as a parametric modulator. The resulting individual contrast images (picture onset > implicit baseline, pleasantness) were compared between groups using independent-samples t-tests on the second level.

In a second first-level model, we examined effects of memory by grouping trial onsets into hit trials (correct old-responses), correct rejection trials (CR, correct new-responses), false alarm trials (FA, incorrect old-responses) and miss trials (cf. Behavioral data). For trials with missing responses (~4% of all trials in both groups), a nuisance regressor was included in the first-level model. Again, pleasantness was modeled as a parametric modulator. The corresponding four individual contrast images (hits > implicit baseline, CR > implicit baseline, FA > implicit baseline, misses > implicit baseline) of interest were fed into a second-level ANOVA. Retrieval success was evaluated by contrasting hits with CR trials using linear contrasts (Spaniol *et al*, 2009) and group differences with regard to retrieval success were evaluated by computing the group by retrieval success interactions (haloperidol hits > CR) > (placebo hits > CR) and (placebo hits > CR) > (haloperidol hits > CR) on the second level. Additionally, we also set up a second-level model including pleasantness per condition.

For the analysis of confidence and to simultaneously examine confidence effects per response category condition, we used a third first-level model where hit, CR, FA and miss trials were further split into high (HC), medium (MC) and low (LC) confidence trials. As before, we included a nuisance regressor for trials with missing responses and modeled pleasantness as a parametric modulator. The corresponding 12 individual contrast images (HC/MC/LC hits > implicit baseline, HC/MC/LC CR > implicit baseline, HC/MC/LC FA > implicit baseline, HC/MC/LC misses > implicit baseline) of interest were fed into a second-level ANOVA. In the second-level analysis, confidence was evaluated by contrasting HC with LC trials using linear contrasts and group differences with regard to confidence were evaluated by computing the group by confidence interactions (haloperidol HC > LC) > (placebo HC > LC) and (placebo HC > LC) > (haloperidol HC > LC). We decided to model confidence as a linear contrast rather than as a parametric modulator of trial onsets because of the group differences present for the trial regressors which might pose a problem for the interpretation of parametric regressor differences.

All resulting activation maps were thresholded at *P* <. 05 (family-wise error (FWE)-corrected for multiple comparisons). Given the strong a-priori hypothesis of involvement of structures with (presynaptic) dopamine receptors, we used a small volume FWE correction (SVC-FWE) at *P* <. 05 based on anatomical masks (50% probability maps) of the striatum and hippocampus created using the Harvard-Oxford cortical and subcortical structural atlases (Desikan *et al*, 2006) as implemented in FSL (https://fsl.fmrib.ox.ac.uk/fsl/fslwiki/Atlases). For all other reported findings, whole-brain FWE-correction at cluster level at *P* <. 05 (using a cluster-forming height threshold at voxel-level of *P* <. 001 (Eklund *et al*, 2016)) was applied. To localize the dopaminergic midbrain (SN/VTA), we used the mean anatomical magnetization transfer (MT) image. The locus coeruleus (LC) was localized using the probability map (2SD map) of the LC (Keren *et al*, 2009) available at http://www.eckertlab.org/LC. Resulting activations in the hippocampus and the amygdala were anatomically localized with probabilistic maps of cytoarchitectonically defined areas (Amunts *et al*, 2005) using the SPM Anatomy toolbox (Eickhoff *et al*, 2005).

## Results

### General drug effects

Comparison of demographic data and all available test data acquired prior to drug administration demonstrated that the groups did not differ with regard to age, sex, weight or in terms of baseline working memory, general response speed or attention capabilities (Table 1). Moreover, participants were not able to guess the substance received above chance level and there were no group differences in reported side effects or subjective feelings relative to baseline (Table S1), confirming the appropriate blinding of participants. With regard to overall group differences, our fMRI results showed significantly higher activity under haloperidol compared to placebo across all recognition trials specifically and selectively in the dorsal striatum (p <. 05 whole-brain FWE-corrected; Fig. 1B). No significant activity differences were found for the inverse contrast (placebo > haloperidol) in any brain region. Importantly, exploration of striatal group differences using an uncorrected threshold did not reveal any striatal voxels showing higher activity for placebo compared to haloperidol.

**Table 1:** Demographics and baseline performance.

|  | Placebo, n = 27 | Haloperidol , n = 27 | Statistics |
| --- | --- | --- | --- |
| Age | 24.37 years ±3.3 | 23.56 years ±2.5 | T(52) = 1.03, p = .31 |
| Sex | 8 males, 19 females | 6 males, 21 females | $\chi^2(1) = 0.39$, p = .54 |
| Weight | 65.70 kg ±6.8 | 65.74 kg ±10.6 | T(52) = -0.15, p = .99 |
| Baseline WM: Complex span/ symmetry accuracy/ RT symmetry rating | 25.78±7.3<br>96.1%±4.2<br>2.17s ±1.44 | 27.85±7.0<br>96.7%±3.7<br>1.67s ±0.63 | T(52) = -1.07, p = .29<br>T(52) = -0.51, p = .61<br>T(35.49) = 1.63, p = .11 |
| Baseline WM: digit span forward/ backward | 8.26±1.3<br>9.15±2.2 | 8.30±1.7<br>8.67±1.7 | T(52) = -0.09, p = .93<br>T(52) = 0.37, p = .93 |
| Baseline WM: block span forward/ backward | 9.73±1.8<br>9.15±1.9 | 10.44±2.0<br>9.81±1.6 | T(51) = -1.35, p = .18<br>T(51) = -1.37, p = .18 |
| Baseline WM: z-summary score | -0.06±0.63 | 0.04±0.58 | T(52) = -0.63, p = .53 |
| Baseline encoding: categorization accuracy picture/ RT/ categorization accuracy arrows/ RT | 76.78%±22.0<br>0.58s±0.04<br>86%±9.0<br>0.38s ±0.01 | 80.23%±15.9<br>0.57s±0.06<br>86%±10.5<br>0.38s ±0.02 | T(51) = -0.66, p = .51<br>T(51) = 0.71, p = .48<br>T(51) = 0.12, p = .91<br>T(51) = 0.05, p = .96 |

± denotes the standard variation

#### Differential effects of haloperidol on recognition performance, metamemory and working memory

Recognition memory performance measured as discrimination sensitivity (d’) computed according to the signal detection theory (SDT) approach (Macmillan and Creelman, 2004) was significantly enhanced in the haloperidol group (t(52) = -2.27, p =. 028, Cohen’s d = -0.63/Pearson’s r = 0.30 (medium effect size); Fig. 2B) which was similarly due to more correct responses for old and new items (i.e., more hits and CR; Table 2) but showed no group difference in response bias (Fig. S1C). Correspondingly enhanced recognition memory under haloperidol was observed when characterizing performance with the corrected hit-rate (Fig. S1B). Moreover, analyzing d’ per confidence level revealed that the superior discrimination performance under haloperidol was driven by better performance on high confidence (HC) trials (significant group×confidence interaction for d’: F(2, 49) = 6.20, p =. 004, partial η^2^ =. 20 (large effect size); post-hoc independent samples t-test HC trials: t(50) = -3.11, p =. 009 (Bonferroni-corrected), Cohen’s d = -0.88/Pearson’s r = 0.40 (large effect size); Fig. 2B). Together, these results indicate that the acute administration of haloperidol boosted retrieval of previously learned information mostly by increasing high confidence correct responses (rather than increasing the amount of correct responses at mid and low confidence range). We additionally analyzed the recognition performance within the framework of the dual processing dual-process signal detection (DPSD) model (Koen *et al*, 2016). Although there was no group×memory-type (recollection/familiarity) interaction (F(1, 52) 0.88, p 0.351), we analyzed the recollection and familiarity effects separately. This explorative analysis revealed that the haloperidol group had significantly higher estimates of recollection only (t(52) = -2.29, p =. 026, Cohen’s d = -0.64/Pearson’s r = 0.30 (medium effect size); Fig. 2C); there was no group difference in the estimates of familiarity (t (52) = -1.43, p = 0.158). However, due to the absence of the interaction effect, one should refrain from drawing specific conclusions from this observation.

**Table 2:**
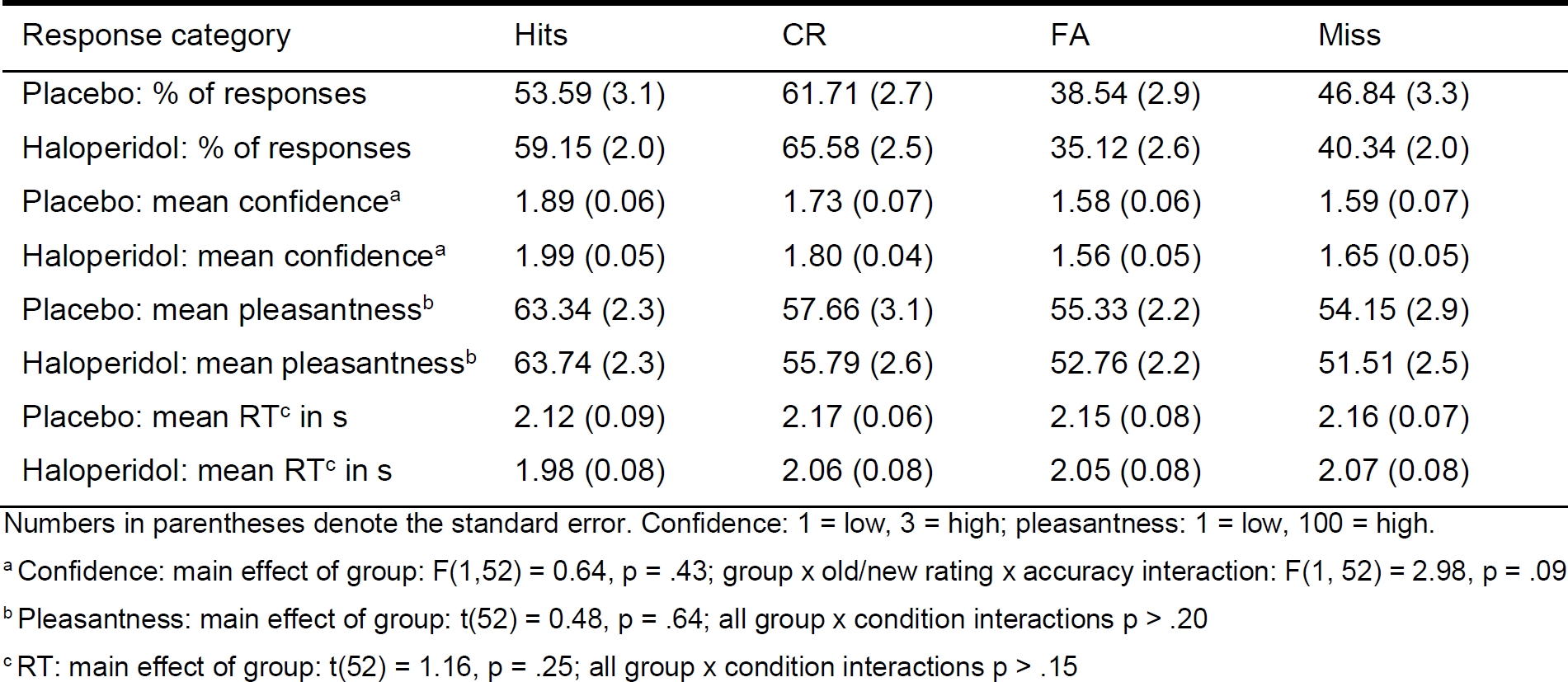
Descriptive statistics of behavioral responses in the recognition memory task.

**Figure 2:**
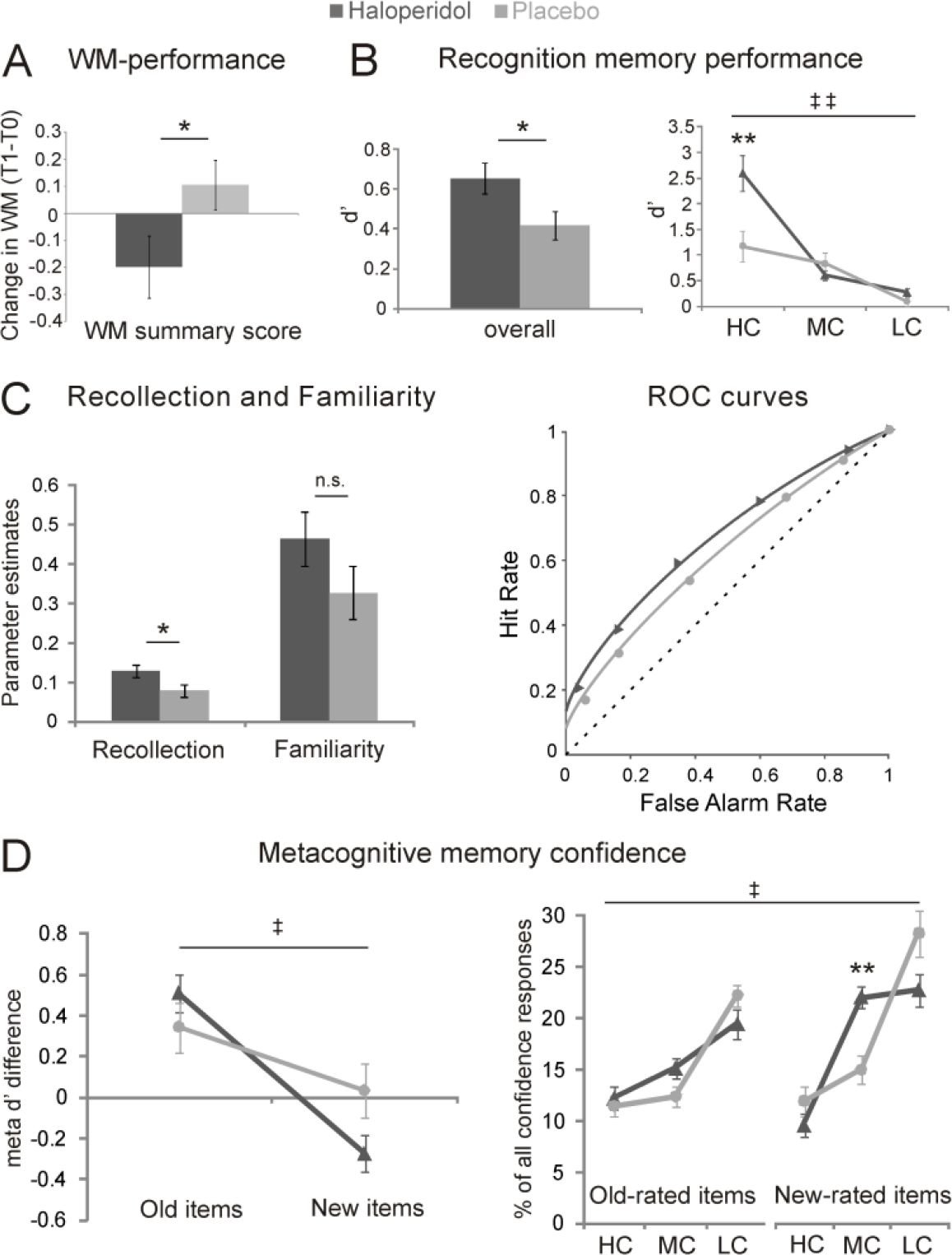
Behavioral effects under haloperidol (dark grey) and placebo (light grey). A) Working memory: Difference in mean z-transformed WM performance after drug administration relative to baseline. B) Memory accuracy: mean d’ overall (left) and d’ per confidence level (right). C) Mean recollection and familiarity parameters (left) and dual-process model-predicted ROC curves (right) with mean observed ROC values (triangles/circles). D) Metacognition: mean meta d’ difference (meta d’ – d’) for old and new items (left) and frequency of confidence ratings for old and new responses (right). */** = significant group difference at p <. 05/at p <. 01 (for post-hoc tests after Bonferroni correction); ǂ /ǂ ǂ = significant group×condition interaction at p <. 05/at p <. 01. n.s. = non-significant. Error bars denote the SEM. HC = high, MC = medium, LC = low confidence.

In contrast, metacognitive memory confidence showed some impairment under haloperidol. We quantified metamemory for old and new picture items using the meta-d’ framework (Maniscalco and Lau, 2012, 2014) which indicates how well the observer’s confidence ratings discriminate between correct and incorrect responses. Importantly, meta d’ is measured in the same units as d’ and therefore it can be compared with d’ to reveal suboptimal (meta-d’ < d’) and enhanced (meta-d’ > d’) metacognition by taking into account the observer’s discrimination sensitivity (Maniscalco and Lau, 2012). Meta d’ difference (meta d’ – d’) showed a significant group×old/new interaction (F(1, 51) = 4.46, p =. 04, partial η^2^ =. 08 (medium effect size)), demonstrating comparably enhanced metamemory for old items in the haloperidol group as in the placebo group but suboptimal metamemory for new items (Fig. 2D). In a similar vein, the frequency of confidence responses showed a significant shift towards medium confidence (MC) responses particularly for new-rated items, indicating reduced confidence differentiation under haloperidol (group×confidence×old/new rating interaction: F(2, 51) = 4.50, p =. 016, partial η^2^ =. 15 (large effect size); post-hoc independent-samples t-test MC new trials: t(52) = -3.44, p =. 006, Cohen’s d = -0.95/Pearson’s r = 0.43 (large effect size) (Bonferroni-corrected); Fig. 2D). As this shift was due to a relative reduction of both HC and LC new-responses, this effect was not reflected by a significant group difference in mean confidence (Table 2; note though that the group×old/new rating×accuracy showed some trend towards significance reflecting the higher mean confidence for hits under haloperidol). Together, these analyses point to an impairing effect of acute haloperidol administration on metacognitive confidence for new-decisions.

There were no group differences for response times or for pleasantness (Table 2 and Fig. S2B). We additionally repeated all above analyses with the baseline WM span summary score (computed from the five behavioral WM span tasks) included as a covariate to control for individual differences in baseline dopamine level (Cools *et al*, 2008). There was no effect on any of the observed group differences, indicating either that baseline dopamine level did not influence the acute haloperidol effects or that the WM span was not a good proxy for baseline dopamine level. Of interest however, there was an detrimental effect of acute haloperidol on WM measured directly after scanning relative to the baseline WM measured before drug administration: the WM summary score demonstrated decreased WM span under haloperidol (time×group interaction: F(1,50) = 4.28, p =. 044, partial η^2^ =. 079 (medium effect size); Fig. 2A and Fig. S1A). However, changes in WM scores relative to baseline did not correlate with the meta-d’ difference for new-decisions (r =. 03, p =. 902) or with memory discrimination performance d’ (r =. 28, p =. 172) across haloperidol participants. Furthermore, there were no significant correlations of weight with memory discrimination performance d’ (r <. 01, p =. 984) or meta-d’ difference for new-decisions (r =. 16, p =. 434) within the haloperidol group. This absence of dosage effects depending on the weight of the participants might not be surprising given the relative restricted range of weight across participants (cf. Table 1).

#### Increased activity in midbrain and mnemonic regions is linked with better recognition performance

Across both groups, the fMRI data revealed a main effect of retrieval success (hits > CR) in various brain regions including the striatum, PFC, hippocampus, substantia nigra/ventral tegmental area (SN/VTA) and anterior cingulate cortex (ACC) (p <. 05 whole-brain FWE-corrected; Fig. 3A). Group differences for retrieval success significant at whole-brain level were evident in an activation cluster localized specifically in the SN/VTA, the hippocampus (presumably CA1 (Amunts *et al*, 2005)), the basolateral amygdala (Amunts *et al*, 2005) and the locus coeruleus (LC) under haloperidol compared to placebo (group×retrieval success interaction, whole-brain FWE-corrected at p <. 05; Fig. 3B and Table 3), suggesting that superior memory performance under haloperidol was linked with higher activity in these midbrain and mnemonic regions. Supporting this role for dopaminergic and hippocampal activity in retrieval performance, the activity in the SN/VTA for retrieval success correlated significantly with memory discrimination (d’) and with right anterior (CA1) hippocampal activity for retrieval success in the haloperidol group (SN/VTA*d’: haloperidol group: r =. 47, p =. 014, placebo group: r =. 16, p =. 445; SN/VTA*hippocampus: haloperidol group: r =. 50, p =. 008, placebo group: r =. 26, p =. 192; Fig. 3B). As these correlations were not significantly different between groups, these correlation findings merely suggest that participants with higher SN/VTA activity showed better recognition memory and had higher hippocampal activity during retrieval in both groups, although this relationship was more pronounced on a descriptive level in the haloperidol group. Of note, we observed no group differences for this retrieval success activity in the striatum even at an uncorrected threshold despite overall increased striatal activity under haloperidol (see also beta plot in Figure 3A).

**Table 3:** Peak activations group differences.

| Region | x | y | z | z-score | Cluster size |
| --- | --- | --- | --- | --- | --- |
| *All trials: H > P* |  |  |  |  |  |
| L caudate nucleus | -12 | 2 | 18 | 4.74 | 348 |
| *Memory effects: Group x retrieval success [H (Hits > CR) > P (Hits > CR)]* |  |  |  |  |  |
| Brain stem | -8 | -36 | -28 | 4.81 | 420 |
| L LC | -6 | -38 | -30 | 3.75 |  |
| R LC | 8 | -38 | -28 | 3.64 |  |
| L SN/VTA | -4 | -20 | -18 | 3.41 |  |
| R Amygdala (LB(Amunts *et al*, 2005)) | 30 | 0 | -24 | 4.02 | 189 |
| R anterior hippocampus (CA1(Amunts *et al*, 2005)) | 30 | -8 | -22 | 3.68 |  |
| R posterior hippocampus (CA2(Amunts *et al*, 2005)) | 34 | -34 | -6 | 3.78<sup>a</sup> | 11 |
| *Confidence effects: Group x confidence [P (HC > LC) > H (HC > LC)]* |  |  |  |  |  |
| L MFG | -26 | 32 | 24 | 5.27 | 2173 |
| L SFG | -22 | 36 | 26 | 5.22 |  |
| R SFG | 16 | 32 | 40 | 4.54 |  |
| R ACC | 6 | 32 | 26 | 4.46 |  |
| L ACC | -8 | 30 | 26 | 4.33 |  |
| R SMA | 10 | 14 | 68 | 4.56 | 230 |
| R NAcc | 10 | 14 | -4 | 4.18<sup>b</sup> | 27 |
| R putamen | 14 | 12 | -6 | 3.93<sup>b</sup> | 23 |
| R caudate nucleus | 12 | 14 | -2 | 3.91<sup>b</sup> | 36 |

x, y, z coordinates refer to the peak voxel in MNI space thresholded at $P < 0.05$ (FWE-corrected at cluster-level; <sup>a</sup>small volume FWE-corrected using the bilateral anatomical hippocampus mask; <sup>b</sup>small volume FWE-corrected using the bilateral anatomical striatum mask). No significant activations were observed for the inverse contrasts. R, right; L, left; H, haloperidol; P, placebo; LC, locus coeruleus; SN/VTA, substantia nigra/ ventral tegmental area; MFG, middle frontal gyrus; SFG, superior frontal gyrus; ACC, anterior cingulate gyrus; SMA, supplementary motor area; NAcc, nucleus accumbens.

**Figure 3:**
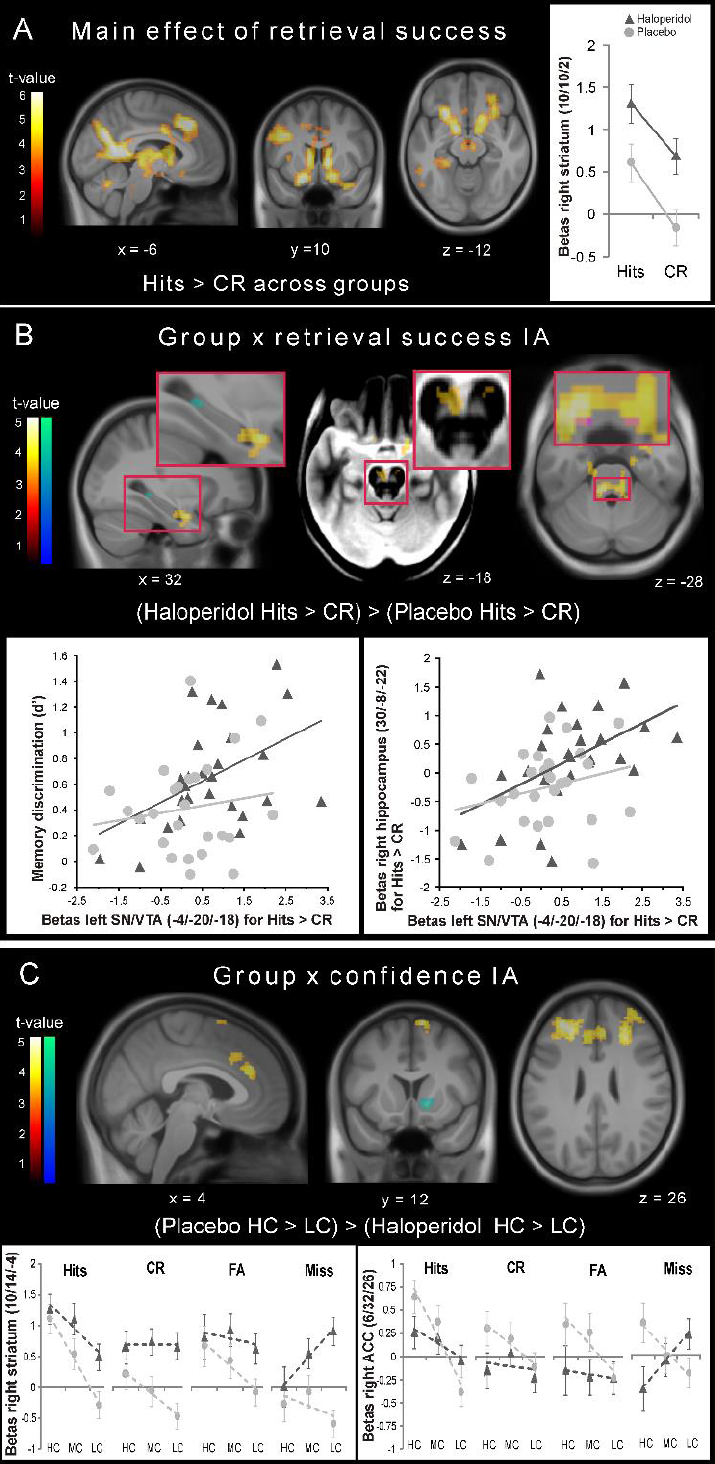
fMRI results. A) Activity pattern of retrieval success across groups and beta estimates from the peak voxel in the right striatum. B) Significantly increased activity for retrieval success under haloperidol in the hippocampus and amygdala (left), SN/VTA (middle; activation displayed on the mean MT image showing the SN/VTA as a bright region) and LC (right; activation overlaid with the LC mask (Keren et al, 2009) in magenta). Scatter plots of the individual SN/VTA response for retrieval success and memory accuracy (left) and the individual anterior hippocampus response for retrieval success (right) in the haloperidol (dark grey triangles) and the placebo group (light grey circles). C) Regions showing higher confidence activity under placebo and beta plots illustrating the confidence pattern in the right striatum and the ACC. Dashed lines represent the linear trend across confidence levels. All activation maps are thresholded at p <. 05 (warm colors: FWE-corrected at cluster-level using a cluster forming threshold at voxel level of p <. 001; cold colors: small volume FWE-corrected using anatomical masks). The inverse contrasts revealed no significant activations. HC = high confidence, MC = medium confidence, LC = low confidence, IA = interaction.

#### Aberrant activity in frontostriatal circuit is linked with impaired metamemory for new-decisions

The main effect of confidence (HC > LC) across groups revealed similar frontostriatal-parietal regions as previous studies (Clos *et al*, 2015; Schwarze *et al*, 2013) (Fig. S2A). The group×confidence interaction showed decreased activity under haloperidol in the PFC and ACC (p <. 05 whole-brain FWE- corrected) and in the right ventral striatum (p <. 05 small volume FWE-corrected; Fig. 3C and Table 3). Importantly, the beta parameters in Figure 3C indicate that this reduced striatal confidence response under haloperidol was due to reduced differentiation between confidence levels rather than due to reduced striatal signal per se: replicating our previous non-pharmacological study (Schwarze *et al*, 2013), the placebo group showed linear decreasing activity in the striatum with decreasing confidence across response category conditions, but this linear confidence effect was clearly reduced under haloperidol due to an unsystematic activity increase across confidence levels. Note that this effect was particularly evident for new-decisions (CR and miss trials). A similar reduced confidence differentiation but without overall increased activity was observed in the ACC. Including the T0 baseline WM span summary score as a covariate in these fMRI analyses did not change the reported group differences for retrieval success or for confidence.

Together, the results show that the dopaminergic modulation improved hippocampally-mediated recognition performance in the haloperidol group but impaired some aspects of frontostriatally- mediated metamemory confidence and working memory.

## Discussion

We showed here that retrieval of previously learned information in humans is under the influence of dopaminergic signaling and that retrieval of episodic information can be improved by acute low-dose D2 antagonist administration. This enhancement of retrieval performance under the D2 antagonist haloperidol is in agreement with previous reports of haloperidol-induced memory retrieval facilitation in the rat (Chugh *et al*, 1991; Sara, 1986). In our current study, the superior retrieval discrimination performance in the haloperidol group was associated with higher activity in the SN/VTA, LC, hippocampus and amygdala compared to the placebo group. Striatal activity was overall increased in this recognition task in the haloperidol group, but there was no evidence for striatal contribution to the improved retrieval performance. Instead, striatal and prefrontal effects were seen for metamemory confidence, where reduced confidence differentiation in the striatum and the PFC mimicked the behavioral impairment in metamemory for new decisions.

While chronic and high-dose administration of haloperidol should decrease dopaminergic signaling due to postsynaptic receptor blockade, acute administration of low doses of D2-antagonists is thought to primarily block presynaptic autoreceptors (which have higher affinity for dopamine than the postsynaptic ones (Ford, 2014)) and thereby lead to paradoxical dopamine-stimulating effects (Frank and O’Reilly, 2006a; Knutson and Gibbs, 2007). Indeed, acute administration of D2-antagonists has been shown to potentiate activity of dopamine neurons and dopamine release in the striatum in response to dopamine-evoking stimuli in rodents and non-human primates (Chen *et al*, 2005; Dugast *et al*, 1997; Garris *et al*, 2003; Jaworski *et al*, 2001; Moghaddam and Bunney, 1990; Pehek, 1999; Pucak and Grace, 1994; Schwerdt *et al*, 2017; Youngren *et al*, 1999). Dopamine-stimulating effects are moreover reflected in increased striatal and hippocampal blood flow in rodents (Chen *et al*, 2005; Schwarz *et al*, 2004) but also in humans (Handley *et al*, 2013) in response to acutely administered D2- antagonists. Generally, the increased striatal and SN/VTA activations in this recognition task under haloperidol seem hard to reconcile with decreased dopaminergic signaling due to postsynaptic receptor blockade, but rather are in accordance with low-dose haloperidol-induced amplification of dopamine release from the SN/VTA. Importantly, optogenetic studies actually demonstrated that phasic stimulation of dopaminergic neurons increases fMRI activity primarily in the dorsal striatum (Ferenczi *et al*, 2016; Lohani *et al*, 2016), which corroborates our dopamine-stimulating interpretation of the haloperidol effects in this recognition task.

D2-autoreceptors are particularly abundant in the striatum (Ford, 2014) but also present in the hippocampus (Rocchetti *et al*, 2015) and amygdala (Bull *et al*, 1991) and although dopamine-stimulating effects in response to D2-antagonists seem to be less extreme in these structures compared to the striatum (Garris and Wightman, 1995), the relative sparseness of DAT in the hippocampus (Kwon *et al*, 2008) might lead to longer-lasting dopamine effects in the hippocampus (Prince *et al*, 2016). In particular, such increased stimulation of the postsynaptic D1-like receptors prevailing in hippocampus and amygdala (Köhler *et al*, 1991; Okubo *et al*, 1999; Rocchetti *et al*, 2015) by the dopaminergic midbrain might have led to the improved memory discrimination, possibly mediated by an increased signal-to-noise ratio (Frank, 2005; Warren *et al*, 2016; Yousif *et al*, 2016) of the representation of episodic information in these mnemonic structures. For example, dopamine might have affected hippocampal pattern completion (Li *et al*, 2010; Wilson *et al*, 2006) or the comparator function of the CA1 region, which compares predicted input computed based on pattern completion in CA3 with actual cortical input during retrieval (Hasselmo *et al*, 1995). Moreover, the strong LC retrieval success activity observed under haloperidol might point to a possible role of noradrenergic signaling in this recognition task (see also (Sara, 1986)). Previous studies reported that acute haloperidol administration leads to increased LC neuron activity possibly by alpha2-adrenergic receptor blockade (Dinan and Aston-Jones, 1984; Olpe *et al*, 1983) and noradrenaline released from the LC into the hippocampus and the amygdala is important for retrieval of emotional memories (Thomas, 2015). Finally, the hypothesis of signal-to-noise ratio amplification is even more prominent for norepinephrine than dopamine (Aston-Jones and Cohen, 2005; Warren *et al*, 2016). However, recent studies revealed that part of the dopaminergic input to the hippocampus actually stems from the dopaminergic co-release of noradrenergic LC neurons (Kempadoo *et al*, 2016; Smith and Greene, 2012; Takeuchi *et al*, 2016). Thus, the strong LC activity for retrieval success might also be due to upregulated dopamine-releasing noradrenergic LC neurons under haloperidol.

In contrast, enhanced dopaminergic stimulation of frontostriatal networks underlying metacognitive (Clos *et al*, 2015) and cognitive control processes (Scimeca and Badre, 2012) under haloperidol resulted in behavioral impairments in metamemory for new-decisions and probably also contributed to the decreased working memory performance (Frank and O’Reilly, 2006b) in the haloperidol group. In particular, the unsystematically increased striatal activity across confidence levels might have limited the dynamic response range of the striatum (Frank, 2005) to signal different confidence levels and altered frontostriatal interactions with the PFC and ACC. These frontostriatal self-monitoring and cognitive control mechanisms seem to be more important for new-decisions than for old-decisions as only the former were impaired under haloperidol. Note that direct stimulating effects of haloperidol in PFC and ACC are less likely due to the relative rareness of presynaptic D2-autoreceptors in these frontal regions (Bannon *et al*, 1983; Lammel *et al*, 2008; Roth, 1984; Wolf and Roth, 1990) and the lack of overall increased activity observed here. However, we can of course not rule out the possibility that the increased striatal activity is caused by reduced inhibition by the PFC under haloperidol rather than the other way around.

In addition to acute haloperidol-induced enhancement of dopamine release, it is possible that postsynaptic effects are present to some degree even at low doses of D2 antagonists. As such, increased dopamine release combined with postsynaptic D2 receptor blockade might decrease the ratio of D2 to D1 receptor activation (Kahnt and Tobler, 2017; Shi *et al*, 1997). Such an activation shift from D2 to D1 receptors might have additionally contributed to the retrieval performance and metamemory effects reported here, rather than increased dopamine release alone.

Of interest, the striatum still showed differential activity for retrieval success in the haloperidol group to a similar degree as in the placebo group. This absence of a striatal retrieval effect under haloperidol might suggest that previously observed retrieval-related activity in the striatum (Clos *et al*, 2015) is not mediated by dopamine. Possibly, this striatal retrieval activity might reflect a glutamatergic signal of hippocampal origin, which is usually thought to signal novelty (Axmacher *et al*, 2010; Lisman and Grace, 2005) but possibly also oldness, depending on the goal of the current state (Herweg *et al*, 2018). However, this null-findings should be treated with some caution as the absence of a striatal retrieval effect might also be due to the lack of power.

Together, these data demonstrate that the dopaminergic modulation here has facilitating effects on midbrain-hippocampal memory retrieval but detrimental effects on frontostriatally-mediated metamemory. These opposing findings are in agreement with recently demonstrated dissociable brain mechanisms underlying recognition performance and memory confidence in non-human primates (Miyamoto *et al*, 2017) and with dissociative effects of noradrenergic modulation on discrimination accuracy and confidence in sensory decision-making (Allen *et al*, 2016; Hauser *et al*, 2017) as well as with long-established behavioral dissociations between memory accuracy and confidence in human cognitive studies (e.g. (Busey *et al*, 2000; Chandler, 1994; Dobbins *et al*, 1998; Roediger and DeSoto, 2014)). Moreover, the improved retrieval performance observed here is consistent with previous rat studies reporting memory facilitation after acute systemic haloperidol administration in the retrieval phase (Chugh *et al*, 1991; Sara, 1986). Finally, our findings might help to explain the presence of episodic memory (Scimeca and Badre, 2012) and metamemory (Eisenacher and Zink, 2017) impairments in dopamine-related diseases as well as offer new perspectives for the treatment of memory disorders.

### Limitations

Our interpretation rests heavily on the assumption that acute administration of 2 mg haloperidol will actually *increase* dopaminergic signaling. However, this effect has only been directly demonstrated in animals. Previous human fMRI studies using similar doses of haloperidol usually assumed that the drug will decrease dopaminergic signaling (e.g., (Oei *et al*, 2012; Pleger *et al*, 2009)), as it does when applied chronically. However, our present fMRI results showing increased activity in striatal and SN/VTA regions under haloperidol seem hard to reconcile with a dopamine-decreasing action of acute low-dose haloperidol in this recognition task. Moreover, it should be noted that these previous findings of decreased striatal signal in response to appetitive stimuli under haloperidol often resulted from contrasting rewarding with non-rewarding stimuli. In the light of the reduced striatal differentiation between confidence conditions (but an overall increased striatal activity) observed in the current study, it *could* be possible that contrasting conditions concealed striatal activity increases under haloperidol in these studies. In other words, reduced striatal differentiation between different conditions might have led to the conclusion of reduced striatal activity, although this activity might effectively have been overall increased (note that contrasting high vs. low confidence in our study resulted in significantly higher activity in the placebo group compared to the haloperidol group). Alternatively, effects of haloperidol might be relatively dependent on the task and the kind of stimuli. Eventually, additional studies including alternative methods such as PET will be necessary to resolve these contradictory drug findings as well to examine the specificity of dopaminergic effects on memory retrieval in more detail. In fact, studies investigating dopaminergic effects on memory retrieval separately from dopaminergic effects on encoding or consolidation are relatively scarce. While two haloperidol studies demonstrated memory facilitation in rats (Chugh *et al*, 1991; Sara, 1986), other animal studies report no (O’Carroll *et al*, 2006; Savalli *et al*, 2015) or, with doses high enough to effectively block postsynaptic D2 receptors, even impairing effects (Blokland *et al*, 1998) of dopaminergic modulation on retrieval discrimination performance. These contradictory results might indicate that the effects might heavily depend on the dopaminergic drug in question, the dosage, the kind of memory task and the route of administration. In humans, the few previous dopamine antagonist memory studies we are aware of modulated dopamine during both encoding and retrieval (Andreou *et al*, 2014; Morcom *et al*, 2010) and therefore are difficult to interpret with regard to retrieval effects specifically.

Moreover, we used a between-group design, which is known to have less power compared to a within-group design. Although we have a reasonable sample size of 27 participants per group and we show strong drug effects both behaviorally and neurally on a whole-brain corrected level, it is possible that some effects were overlooked due to lack of power. In particular the absence of the striatal retrieval effect might be due to too little power and should therefore be interpreted with caution.

## Author contributions

Conceptualization, MC and TS; Methodology, MC and TS; Formal Analysis, MC; Investigation, MC; Writing – Original Draft, MC; Writing – Review & Editing, MC, NB and TS; Resources – NB and TS; Supervison – TS; Funding Acquisition, NB and TS.

## Acknowledgments

This study was supported by a grant from the Deutsche Forschungsgemeinschaft to TS and NB (DFG SO 952/3-1). The funders had no role in study design, data collection and analysis, decision to publish, or preparation of the manuscript. The authors are grateful to the University of Minnesota Center for Magnetic Resonance Research for providing the image reconstruction algorithm for the simultaneous multi-slice acquisitions.

